# *Haemoproteus paraortalidum* n. sp. in captive Black-fronted Piping-guans *Aburria jacutinga* (Galliformes, Cracidae): high prevalence in a population reintroduced into the wild

**DOI:** 10.1101/280990

**Authors:** Francisco C. Ferreira-Junior, Daniela de Angeli Dutra, Nelson R.S Martins, Gediminas Valkiūnas, Érika M. Braga

**Author notes:** Corresponding authors (FCFJ); (EMB). Departamento de Parasitologia, Universidade Federal de Minas Gerais, Avenida Pres. Antônio Carlos 6627, Belo Horizonte, MG, 31270-901, Brazil.

## Abstract

Haemosporidian parasites of the genus *Haemoproteus* are widespread and can cause disease and even mortality in birds under natural and captive conditions. The Black-fronted Piping-guan (*Aburria jacutinga*) is an endangered Neotropical bird of the Cracidae (Galliformes) going through a reintroduction program to avoid extinction. We used microscopic examination and partial cytochrome *b* DNA sequencing to describe a new *Haemoproteus* species infecting Black-fronted Piping-guans bred and raised in captivity that were reintroduced into the Atlantic rainforest. *Haemoproteus* (*Parahaemoproteus*) *paraortalidum* n. sp. was detected in the blood of 19 out of 29 examined birds. The new species is distinguished from other haemoproteids due to the shape of gametocytes, which have pointed ends in young stages, and due to the presence of vacuole-like unstained spaces in macrogametocytes and numerous volutin granules both in macro- and microgametocytes. Illustrations of the new species are provided. Phylogenetic inference positioned this parasite in the *Parahaemoproteus* subgenus clade together with the other two *Haemoproteus* genetic lineages detected in cracids up to date. We discuss possible implications of the reintroduction of birds infected with haemosporidian parasites into de wild. Treatment of *Haemoproteus* infections remains insufficiently studied, but should be considered for infected birds before reintroduction to improve host reproductive and survival rates after release.

## 1. Introduction

Birds of the family Cracidae (order Galliformes) are endemic to the Neotropical forests of Central and South America. With 19 out of 50 species listed as vulnerable, endangered or critically endangered, this is the most threatened avian family in the continent (Pereira et al., 2000). The Black-fronted Piping-guan (*Aburria jacutinga*) is an endangered medium sized (1.1-1.4 Kg) cracid from the Brazilian Atlantic rainforest facing local population declines and extinctions due to habitat loss and illegal hunting (Bernardo et al., 2011b; ICMBio, 2008). This bird species is one of the most important seed dispersers from the Atlantic rainforest (Galetti et al., 2013), playing a crucial role in this highly endangered biodiversity hotspot (Myers et al., 2000). Currently, five conservation centrers breed this species for reintroduction under the Brazilian Action Plan for the Conservation of Endangered Galliformes (ICMBio, 2008; Oliveira-Jr. et al., 2016). Reintroduction of captive-bred animals is essential to avoid threatened birds from being extinct (Costa et al., 2017; Earnhardt et al., 2014; Hammer and Watson, 2012) and is successful for the Red-billed Curassow (*Crax blumenbachii*), for example, a Brazilian cracid that has been reintroduced in the Atlantic rainforest for almost 30 years (Bernardo et al., 2011a; São Bernardo et al., 2014).

Infectious diseases may compromise such programs by reducing longevity and reproduction of captive animals (Sainsbury and Vaughan-Higgins, 2012) and by excluding animals from reintroduction (Ewen et al., 2012). To comply with international guidelines (IUCN, 2009), health assessments were conducted to identify hazards to captive populations of Black-fronted Piping-guans participating in reintroduction programs, revealing infections by mites, helminths and haemosporidian parasites (Marques et al., 2013; Motta et al., 2013).

Avian haemosporidians are widely distributed protozoans belonging to the genera *Plasmodium, Haemoproteus, Leucocytozoon* and *Fallisia*, and infect most bird groups in all continents except Antarctica (Clark et al., 2014; Valkiūnas, 2005). Haemosporidian prevalence vary between different bird species managed under captive conditions, and *Haemoproteus* parasites, for instance, are associated with mortality episodes in a wide range of species (Cardona et al., 2002; Donovan et al., 2008; Olias et al., 2011; Pacheco et al., 2011; Valkiūnas and Iezhova, 2017). Consequently, these infections can be detrimental to *ex situ* populations, and can hamper efforts to supply birds to reintroduction programs. Moreover, haemosporidians can be accidently introduced and established into the wild together with their hosts, with unpredictable consequences for local avian community (Beadell et al., 2006; Castle and Christensen, 1990).

Two *Haemoproteus* species have been described at morphospecies level in Cracidae birds, *Haemoproteus cracidarum* (Bennett et al., 1982) and *Haemoproteus ortalidum* (Gabaldon and Ulloa, 1978). Molecular characterization is available only for *H. ortalidum* (Chagas et al., 2017), demonstrating a likely underestimation of *Haemoproteus* parasites infecting birds of this Neotropical bird family. Moreover, these two parasite species together with *Haemoproteus mansoni* (synonym of *Haemoproteus meleagridis*), *Haemoproteus lophortyx* and *Haemoproteus stableri* constitute the entire known diversity of *Haemoproteus* parasites infecting birds of Galliformes in the American continent (reviewed in Valkiūnas, 2005). Here, we describe a new *Haemoproteus* (*Parahaemoproteus*) species from a captive population of Black-fronted Piping-guan. Because these birds have been bred in captivity and reintroduced into the Atlantic rainforest in southeastern Brazil, we also pointed possible implications related to the reintroduction of birds infected with haemosporidian parasites to the wildlife.

## 2. Material and methods

### 2.1. Captive population and sample collection

We analysed the presence of haemosporidian parasites in Black-fronted Piping-guan individuals hatched and raised in CRAX Brazil — Wildlife Research Society, a conservation centre situated in Contagem municipality, Minas Gerais state, Brazil (19 °51’05”S, 44 °04’03”W). This facility focuses on the reproduction of endangered birds for reintroduction programs, and possesses approximately 1500 birds of 50 species, mainly belonging to Accipitriformes, Columbiformes, Galliformes and Tinamiformes. This conservation centre manages Black-fronted Piping-guans in breeding pairs in separate enclosures, as well as small groups of young and adult birds in collective aviaries. All enclosures are fenced with 50 mm wire mesh and birds receive manufactured ration with no antibiotics or anticoccidials once daily. Water is provided *ad libitum*. The reintroduction area consists of a 560 ha fragment of secondary Atlantic forest in Ipaba municipality (19 °26’26”S, 42 °11’45”W), Brazil, 180 km from CRAX Brazil.

We sampled 29 birds between September 2013 and March 2014 in CRAX Brazil on the day these birds were transported to the release site. Before being released into the wild, birds remained in a meshed enclosure at the border of a protected secondary forest for approximately 40 days, where they received food and water *ad libitum*. After this acclimatization period, birds were released and the enclosure gates are kept opened so that birds have free access to these provisions.

We obtained blood samples (∼ 100 µL) through brachial venipuncture. Two blood films were prepared from each bird, and the material was rapidly air dried and subsequently fixed with absolute methanol for one minute within six hours after preparation. We stored the remaining volume of blood samples in absolute ethanol at -20°C until DNA was Çextracted. This study was approved by the Ethics Committee in Animal Experimentation (CETEA), Universidade Federal de Minas Gerais, Brazil (Protocol #254/2011) and by Brazilian environment authorities (SISBIO 16359-3).

### 2.2. Microscopic examination

Blood films were stained with 10% Giemsa solution (pH = 7.2) for 40 minutes within two days of preparation. An Olympus CX31 light microscope equipped with an Olympus Q-Color5 imaging system (Olympus, Tokyo, Japan) and Q-Capture Pro7 imaging software (QImaging, Surrey, Canada) were used to examine blood films and to capture images from all sampled birds. At least 200 microscopic fields under 1000 × magnification were examined for the detection of parasites. Morphometric features studies (Table 1) were those defined by Valkiūnas (2005), and two-tailed Student’s *t*- test for independent samples was used to determine whether parasites affect the morphology of their host cells. For this, we compared the length and width of infected and uninfected cells. A *P-*value of 0.05 or less was considered significant. Parasitemia intensity was estimated by actual counting the number of infected erythrocytes per 20,000 observed erythrocytes.

**Table 1.** Morphometry of host cells and mature gametocytes of *Haemoproteus* (*Parahaemoproteus*) *paraortalidum* n. sp. from Black-fronted Piping-guan *Aburria jacutinga*.

| Feature | Measurements (μm)<sup>a</sup> |
| --- | --- |
| **Uninfected erythrocyte** | **N = 15** |
| Length | 11.4 – 14.3 (12.7 ± 0.63) |
| Width | 6.9 – 8.5 (7.7 ± 0.46) |
| **Uninfected erythrocyte nucleus** | **N = 15** |
| Length | 5.4 – 7.4 (6.2 ± 0.6) |
| Width | 2.4 – 3.1 (2.7 ± 0.2) |
| **Macrogametocyte** | **N = 6** |
| **Infected erythrocyte** |  |
| Length | 10.7 – 13.4 (12 ± 0.81) |
| Width | 6.6 – 9.1 (8.1 ± 0.95) |
| **Infected Erythrocyte Nucleus** |  |
| Length | 5.1 – 6.6 (5.76 ± 0.53) |
| Width | 2.3 – 3.0 (2.61 ± 0.25) |
| **Gametocyte** |  |
| Length | 9.4 – 12.1 (10.97 ± 1.12) |
| Width | 3.7 – 5.9 (4.23 ± 0.75) |
| **Gametocyte nucleus** |  |
| Length | 2.3 – 2.9 (2.54 ± 0.18) |
| Width | 1.4 – 2.3 (1.81 ± 0.32) |
| Number of pigment granules | 10.0 – 19.0 (14.33 ± 2.98) |
| Diameter of pigment granules | 0.5 – 1.2 (0.68 ± 0.13) |
| Vacuole | 1.6 – 2.0 ($1.76 \pm 0.13$) |
| NDR | 0 – 0.52 ($0.33 \pm 0.19$) |
| **Microgametocyte** | **N = 4** |
| **Infected erythrocyte** |  |
| Length | 11.7 – 13.4 ($12.68 \pm 0.77$) |
| Width | 6.5 – 8.4 ($7.63 \pm 0.88$) |
| **Infected Erythrocyte Nucleus** |  |
| Length | 4.6 – 6.2 ($5.29 \pm 0.65$) |
| Width | 2.2 – 2.7 ($2.5 \pm 0.28$) |
| **Gametocyte** |  |
| Length | 10.7 – 12.6 ($11.98 \pm 0.85$) |
| Width | 2.9 – 3.5 ($3.24 \pm 0.24$) |
| **Gametocyte nucleus** |  |
| Length | 4.1 – 10.2 ($7.16 \pm 2.6$) |
| Width | 2.2 – 3.1 ($2.59 \pm 0.36$) |
| Number of pigment granules | 9 – 16 ($12.7 \pm 2.87$) |
| Diameter of pigment granules (N = 27) | 0.5 – 1.0 ($0.65 \pm 0.13$) |
| NDR | 0.3 – 0.8 ($0.61 \pm 0.2$) |

<sup>a</sup> All measurements are given in micrometers except for the items “Number of pigment granules”. Minimum and maximum values are provided followed in parentheses by the arithmetic mean and standard deviation.
NDR = nucleus displacement ratio according to Valkiūnas (2005).

### 2.3. Molecular and phylogenetic analyses

Approximately 10 µL of blood was transferred to 1.5 mL microtubes and samples were dried at 37 °C for subsequent DNA extraction, for which we used a conventional phenol-chloroform method with isopropanol precipitation (Sambrook and Russell, 2001). The genomic DNA pellet was resuspended in 50 µL of ultrapure water and quantified using a NanoDrop 2000 (Thermo Scientific, Waltham, United States). Between 50 and 100 ng of the extracted DNA was used for a screening PCR that amplifies a 154-nucleotide segment (excluding primers) of ribosomal RNA coding sequence within the mitochondrial DNA of *Plasmodium* and *Haemoproteus* in a single reaction. We used the primers 343F (5’-GCTCACGCATCGCTTCT-3’) and 496R (5’-GACCGGTCATTTTCTTTG-3’) designed by Fallon et al (2003) under PCR conditions and amplification analysis described by Roos et al (2015). We used Fisher’s exact test to compare haemosporidian detection between microscopic examination and PCR.

DNAs from positive individuals were submitted to a nested-PCR targeting the amplification of a 478 bp region of the cytochrome b gene (*cytb*). For the first reaction, we used primers HaemNFI (5’-AGACATGAAATATTATGGITAAG-3’) and HaemNR3 (5’-GAAATAAGATAAGAAATACCATTC-3’) (Hellgren et al., 2004) with 50-100 ng of genomic DNA. A 1-μL aliquot of this PCR product was then used as a template for the second reaction with the primers HaemF (5’-CTTATGGTGTCGATATATGCATG-3’) and HaemR2 (5’-CGCTTATCTGGAGATTGTAATGGTT-3’) (Bensch et al., 2000). Both reactions contained 1× buffer, 4 mM of MgCl2, 0.3 mM of each dNTP, 1 unit of Taq (Phoneutria, Belo Horizonte, Brazil), 0.4 mM of each primer, and nuclease-free water in 25μl reaction volumes. DNA extracted from blood samples of chickens experimentally infected with *Plasmodium gallinaceum* and ultrapure water was used as positive and negative controls, respectively. These nested-PCRs followed the protocol by Hellgren et al (2004).

Products from all positive nested-PCRs were purified with Polyethylene Glycol 8000 (Sambrook and Russell, 2001) and bi-directionally sequenced with dye-terminator fluorescent labeling in an ABI Prism 3100 sequencer (Applied Biosystems, Foster City, United States). DNA sequences were aligned, checked for the presence of mixed infections (presence of double peaks in the electropherogram, edited using ChromasPro 2.0.6 (Technelysium Pty Ltd, Helensvale, Australia), and compared with data available in the public databases Genbank and MalAvi. Detected *cytb* sequence was deposited in GenBank under accession number MH036944.

A Bayesian phylogenetic reconstruction was performed. We compared the parasite sequence found in this study with haemoproteids of subgenera *Parahaemoproteus* and *Haemoproteus* found in birds of Galliformes and in other orders. This inference was produced using MrBayes 3.2.2 (Ronquist and Huelsenbeck, 2003) with the GTR + I + G model of nucleotide evolution, as recommended by ModelTest (Posada and Crandall, 1998), which selects the best-fit nucleotide substitution model for a set of genetic sequences. We ran two Markov chains simultaneously for 5 million generations in total that were sampled every 1000 generations. The first 1250 trees (25%) were discarded as a burn-in step and the remaining trees were used to calculate the posterior probabilities of each estimated node in the final consensus tree. We used *Leucocytozoon caulleryi* as the outgroup to root the phylogenetic tree.

## 3. Results

### 3.1. Prevalence data

The screening PCR detected 18 positive birds (62.1%). Microscopic examination of blood films demonstrated the presence of haemosporidians in 13 samples (44.8%), with parasitemia ranging between 0.005% and 0.1%. Detection of parasites was similar between the microscopic examination and PCR (*P* = 0.29). Only one sample positive in microscopic examination was negative in the screening PCR (parasitemia = 0.005%), and six samples positive in this PCR were negative by microscopic examination. We obtained 18 positive samples in the *cytb* PCR, and high-quality sequences were obtained from 15 samples, revealing identical parasite lineage (GenBank accession number MH036944). Both microscopic examination and PCR-based detection revealed single *Haemoproteus* infections in all positive birds.

### 3.2. Description

***Haemoproteus* (*Parahaemoproteus*) *paraortalidum*** n. sp. (Fig. 1, Table 1) *Young gametocytes* (Fig. 1A-F): Develop in mature erythrocytes and usually present in a position ranging from lateral-polar (Fig. 1A, C) to lateral (Fig. 1B) to nuclei of the host cells. Earliest gametocytes are oval in shape (Fig. 1A, B); growing young gametocytes are slender rod-like bodies, with even outline and often with slightly pointed ends (Fig. 1C); they lie free in the cytoplasm and do not touch either the nucleus or envelope of erythrocytes (Fig. 1A-C). Nuclei are dark-stained, prominent, usually even in outline, with well-distinct boundaries, features that are particularly visible in early gametocytes (Fig. 1B, C). Nuclei assume various positions in early gametocytes, but were more often seen close to the central or sub-central positions in advanced young gametocytes (Fig. D-F). As parasite grows, gametocytes extend along the erythrocyte nuclei; they assume irregular elongate shapes, with ends thinner than the gametocyte width (Fig. 1F). In this stage of development, gametocytes usually lie free in the cytoplasm, but might also touch the nuclei or envelope of erythrocytes occasionally. A single small vacuole appears in advanced young gametocytes; it locates close to parasite nucleus (Fig. 1D, E). Pigment granules are blackish-brown, of small (< 0.5 µm) size, often grouped. Volutin granules appear as tiny spots in early gametocytes, and volutin amount increases as parasites grow. Volutin tend to gather along the periphery of gametocytes being particularly dense on gametocyte tips, and as a result of this, central parts of advance gametocytes look paler-stained than their peripheral parts (Fig. 1D-F). The influence of young gametocytes on infected erythrocytes is not pronounced, a characteristic feature of this species development.

**Fig. 1.**
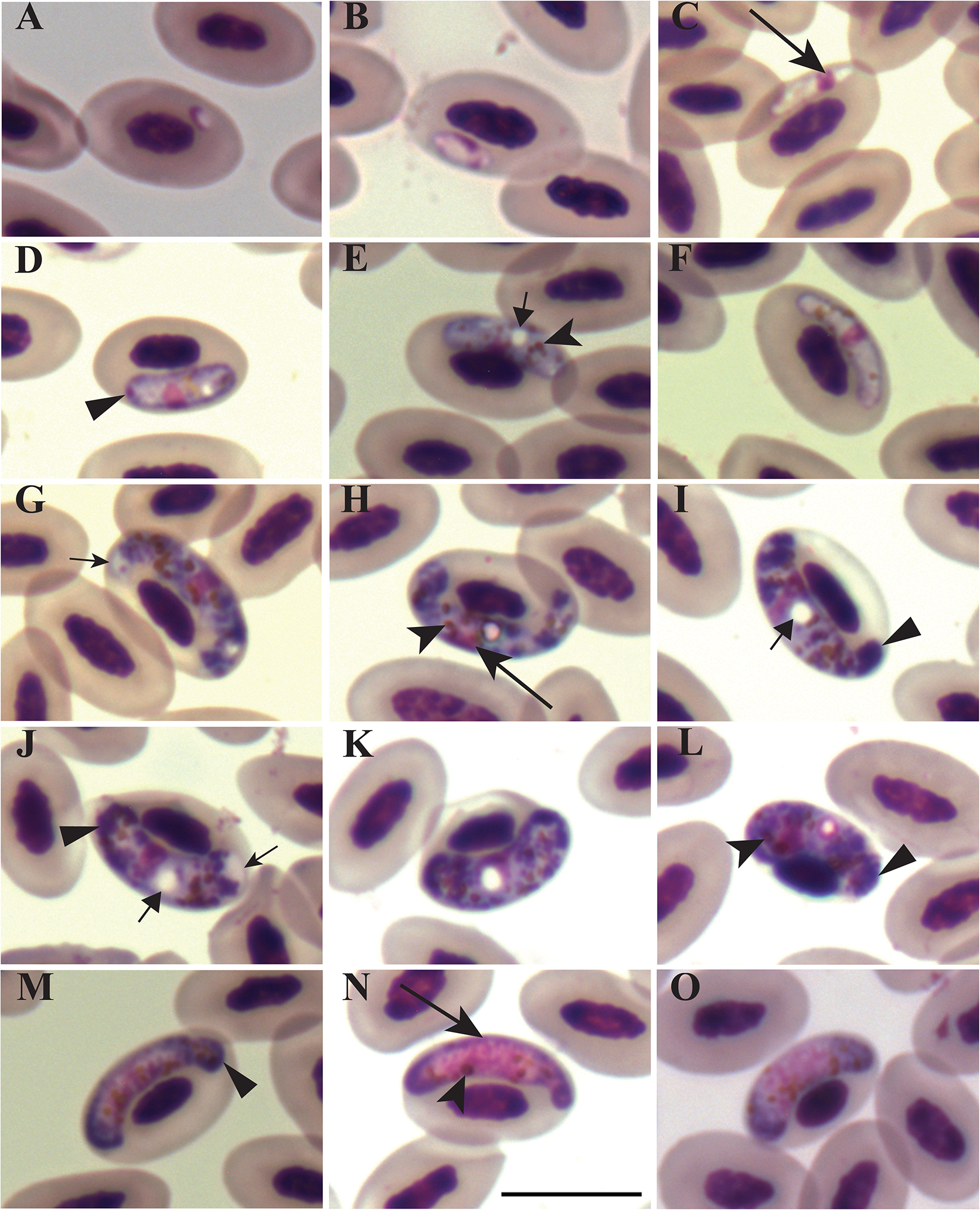
*Haemoproteus* (*Parahaemoproteus*) *paraortalidum* n. sp. from Black-fronted Piping-guans *Aburria jacutinga*. A–F: young gametocytes. G–L: macrogametocytes. M–O: microgametocytes. Long simple arrows - nuclei of parasites. Short simple arrows - vacuole-like unstained spaces on the ends of gametocytes. Simple arrowheads - pigment granules. Short triangle arrows - vacuoles. Triangle arrowheads - clumps of volutin granules. Giemsa stained thin blood films. Scale bar = 10 μm.

*Macrogametocytes* (Fig. 1G-L): The cytoplasm is granular in appearance, contains numerous volutin granules, which tend to group along the periphery of gametocytes, being particularly densely aggregated on the ends of gametocytes (Fig. 1H-J), a characteristic feature of this species development. The volutin granules are similar to pigment granules in size and shape, often grouped, and they obscure the pigment granules. Gametocytes grow along the nuclei of infected erythrocytes (Fig. 1G-L). Advanced growing parasites adhere to the envelop of erythrocytes, but usually do not touch nuclei of erythrocytes and, as a result of this, a more or less evident unfilled space (a ‘cleft’) frequently occurs between gametocytes and the nuclei of erythrocytes (Fig. 1G-I). In fully grown gametocytes, this space might be evident, but not in all cells (Fig. 1J, K). Ends of growing gametocytes are markedly thinner than the central part of gametocytes (Fig. 1G-I), and they enclose slightly the erythrocyte nuclei giving a ‘horn’-like appearance to the parasite due to more or less symmetrical position of the ends in regard to the nuclei of erythrocytes (Fig. 1H). Ends of maturing gametocytes are slender, wavy or slightly ameboid in outline (Fig. G-I); they slightly enclose nuclei of erythrocytes, but do not fill the polar region of host cells. Maturing gametocytes are broad sausage-like bodies, with even outline (Fig. 1J-L); they displace host cell nuclei laterally, sometimes markedly (Fig. 1L). Vacuole-like unstained spaces were visible on one or both ends in about 37% of growing and mature gametocytes (Fig. 1G, J), a characteristic feature of this species development. Parasite nuclei are compact, relatively small (Table. 1), of variable shape, usually of central (Fig. 1G) or subcentral (Fig. 1H-J) position. One large (>1 µm), readily visible vacuole is present in each macrogametocyte (Fig. 1I-J), a characteristic feature of this species development. Such vacuoles usually are closely associated with gametocyte nuclei (Fig. 1H-K) or locate close to the nuclei (Fig. 1L). Pigment granules are roundish or oval, of medium size (between 0.5 and 1 μm in length), scattered throughout the cytoplasm or grouped The influence of growing gametocytes on host cell is not pronounced (Fig. 1G-I). Fully grown gametocytes markedly displace host cell nucleus laterally (Fig. 1J-L) however, no significant differences were reported in the length (*P* = 0.12) or width (*P* = 0.43) of infected cells in comparison to uninfected ones (Table 1).

*Microgametocytes* (Fig. 1M-O): The general configuration and other features are as for macrogametocytes with the usual haemosporidian sexual dimorphic characters, which are the pale staining of the cytoplasm and diffuse pale-stained nuclei. The proportion of macro- and microgametocytes was 4: 1 in the type material. Few non-deformed microgametocytes were seen and measured (Table 1). Vacuoles in the cytoplasm or vacuole-like spaces on the ends of parasites were not seen at any stage of microgametocyte development.

### 3.3. Taxonomic summary

Type host: Black-fronted Piping-guan *Aburria jacutinga* (Spix, 1825) (Galliformes, Cracidae).

DNA sequences: Mitochondrial cyt *b* lineage ABUJAC01 (478 base pairs) with GenBank accession MH036944.

Type locality: Contagem, Minas Gerais State, CRAX Brazil— Wildlife Research Society, Brazil (19 °51’05”S, 44 °04’03”W).

Type specimens: Hapantotypes (accession number LABMAL001 and LABMAL002, parasitemia intensities are approximately 0.04% and 0.1%, respectively, lineage ABUJAC01, *Aburria jacutinga* collected by F. Ferreira-Junior, Contagem municipality, Brazil 16 September 2013) and parahapantotypes (accession numbers 49019 NS and 49020 NS, duplicate of the hapantotype) were deposited in the Institute of Biological Sciences (Belo Horizonte, Brazil) and in Nature Research Centre (Vilnius, Lithuania), respectively.

Site of infection: Mature erythrocytes; no other data.

Prevalence: Nineteen of 29 investigated Black-fronted Piping-guan (65.5%) were positive (screening PCR and microscopic examination combined data).

Additional hosts and distribution: This parasite lineage was detected in Black-fronted Piping-guans from the same conservation centre in 2010 (GenBank GenBank accession KC250002, Motta et al., 2013). The same lineage was also detected in two Yellow-breasted Flycatchers (*Tolmomyias flaviventris*) captured in Aracruz, Espírito Santo State, Brazil (GenBank accession JX029916; Lacorte et al., 2013).

Etymology: The species name reflects some similarities of morphological features of this parasite gametocytes to those of *Haemoproteus ortalidum*, particularly presence of one large circular vacuole in each macrogametocyte. Additionally, the new species and *H. ortalidum* parasitize birds of the Cracidae.

### 3.4. Remarks

The new species share some morphological characteristics with *Haemoproteus ortalidum*, which was found in the Rufous-vented chachalaca *Ortalis ruficauda* in Venezuela (Bennett et al., 1982; Gabaldon and Ulloa, 1978), as both parasites have halteridial macrogametocytes, each possessing one large vacuole. The new species can be readily distinguished from *H. ortalidum* primarily due to 1) the presence of numerous volutin granules, which overfill mature macrogametocytes (Fig. 1G-O), 2) the presence of growing gametocytes that have ends markedly thinner than the central part of the gametocytes (Fig. 1G-I), and 3) the presence of vacuole-like unstained spaces on one or both ends of maturing and fully grown macrogametocytes (Fig. 1G, J). These are not characters of *H. ortalidum*. Additionally, cylindrical form of growing gametocytes and roundish (discoid) form in mature gametocytes are common in *H. ortalidum*, but are not characteristic of the new species.

### 3.5. Phylogenetic relationship of parasites

Our phylogenetic analysis showed a well-supported clade formed by all *Haemoproteus* detected in cracids up to date. For this analysis, we included a parasite lineage detected in White-browed Guan (*Penelope jacucaca*) sampled in Peru (unpublished work; GenBank accession KF482345). This parasite have 1% of genetic divergence (polymorphisms in five nucleotides out of 478) when compared to *H. paraortalidum* n. sp. (Fig 2). Additionally, *H. ortalidum*, the only *Parahaemoproteus* parasite infecting cracids and characterised on molecular and morphological levels (Chagas et al., 2017), cluster in the same clade (Fig. 2). These parasite morphospecies differ at 4.2% in the *cytb* gene (polymorphisms in 18 nucleotides out of 478). This genetic difference in *cytb* gene is above the divergence threshold at which two parasite lineages are expected to be morphologically different (Hellgren et al., 2007), showing accordance between molecular and morphological characterization of *Parahaemoproteus* parasites infecting cracids.

**Fig. 2.**
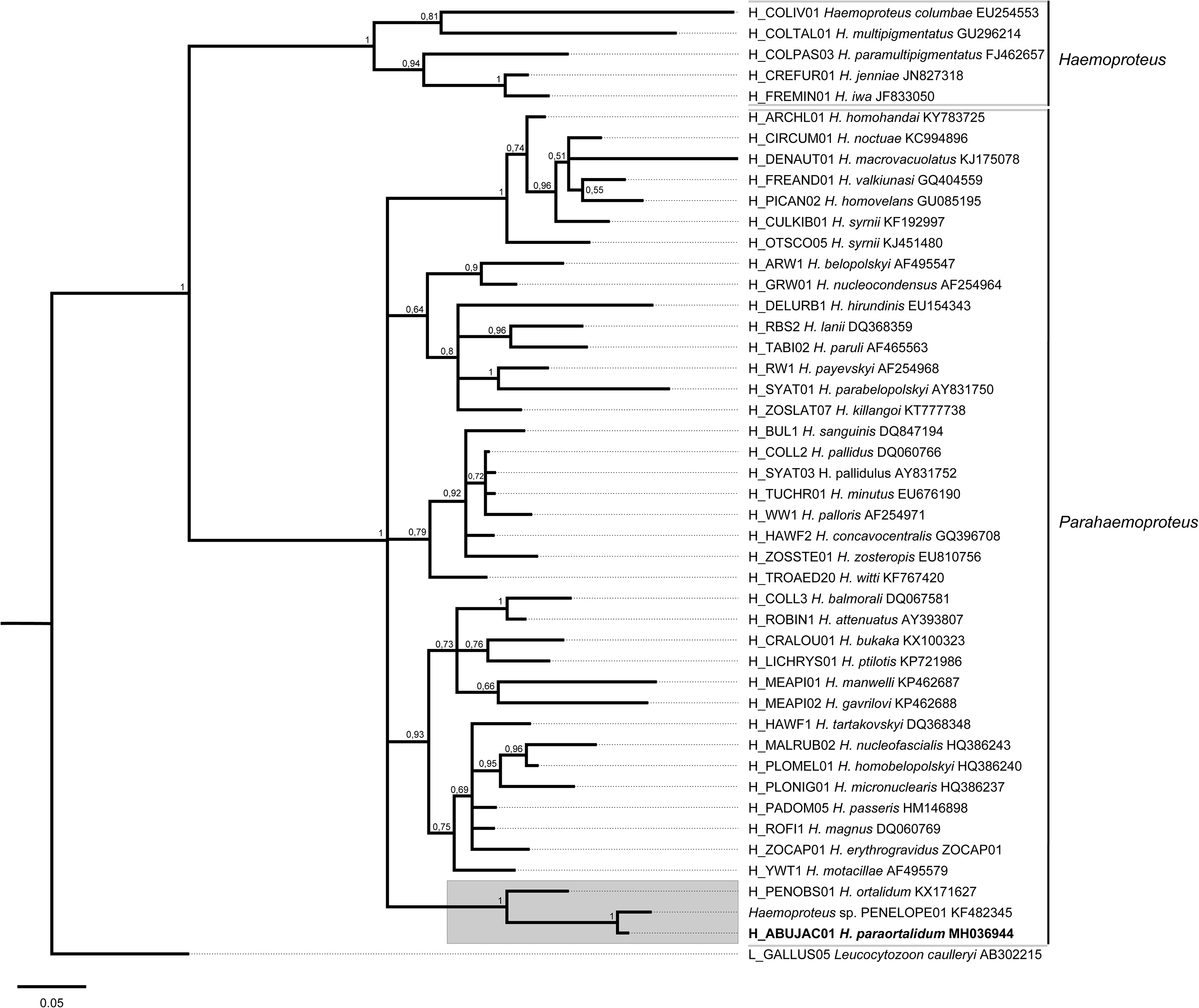
Bayesian phylogenetic tree showing the relationships between *Haemoproteus* (*Parahaemoproteus*) *paraortalidum* n. sp. from Black-fronted Piping-guans *Aburria jacutinga* (in bold) and related parasites of subgenera *Parahaemoproteus* and *Haemoproteus*. *Haemoproteus* parasites detected in cracid birds are indicated by a grey box. Values near branches are posterior probabilities. A sequence of *Leucocytozoon caulleryi* was used as outgroup.

## 4. Discussion

Avian haemosporidians, together with other agents of infectious diseases, can constitute an important threat to wild birds under *in situ* and *ex situ* conditions. Here, we described a new blood parasite infecting a captive population of Black-fronted Piping-guan reintroduced into the wild. This is the first morphological and molecular characterization of *Haemoproteus* species reported in Black-fronted Piping-guan, an endangered bird under rapid population decline. Avian haemoproteids are transmitted by louse flies of the Hippoboscidae (parasites of subgenus *Haemoproteus*) and biting midges of the Ceratopogonidae (parasites of subgenus *Parahaemoproteus*) (Valkiūnas, 2005). Phylogenetic analysis placed *Haemoproteus paraortalidum* n. sp. in a clade of parasites belonging to the *Parahaemoproteus* subgenus (Fig. 2), which numerous species are transmitted by biting midges (Diptera, Ceratopogonidae: *Culicoides* spp.) (Bukauskaitė et al., 2015). Hippoboscid-transmitted species of subgenus *Haemoproteus* appeared in a separate well-supported sister clade. Recent experimental studies show that phylogenies based on *cytb* gene indicate vectors of avian haemoproteids (Bukauskaitė et al., 2015; Žiegytė et al., 2017), providing opportunities to identify possible vectors of haemoproteid species.

We detected high prevalence (65.5%) of *H. paraortalidum* n. sp. in individuals that were born and raised in captivity, showing active local transmission of this infection by vectors that are likely to be biting midges of genus *Culicoides*. Black-fronted Piping-guan were maintained in enclosures covered by nets with large mesh size that allowed free movement of biting midges between bird groups. To protect birds from *Haemoproteus* infections we recommend growing of chicks, which are particularly vulnerable to various haemosporidian infections (Cardona et al., 2002; Graczyk et al., 1994), in closed aviaries protected from penetration of biting midges.

A previous study at CRAX Brazil reported *Haemoproteus* infection in 42.8% (n=21) in Black-fronted Piping-guan, with the detection of the same *H. paraortalidum* n. sp. lineage described here (Motta et al., 2013). However, there were no significant alterations in hematological, biochemical and in the serum protein (albumin and immunoglobulins) profiles due to this infection, indicating that this parasite may not cause major deleterious effects under the same captive circumstances as evaluated here. Birds in our study and in Motta et al. (2013) showed low parasitemia (ranging from undetectable parasites under microscopy to 0.1%), a characteristic of the chronic stage of infection (Valkiūnas, 2005). These birds were provided with food and water, and were protected from harsh environmental conditions and predators. This might facilitate survival and recovery during acute stage of *Haemoproteus* infection, at which hosts suffer from physiological alterations (Valkiŭnas et al., 2006). Indeed, galliform birds can survive infection of pathogenic *Haemoproteus* parasites under experimental (Atkinson et al., 1988) and captive (Cardona et al., 2002) conditions after presenting clinical signs of disease at levels that would likely reduce survival rates in the wild. These aspects warrant further studies to evaluate the pathogenicity of *H. paraortalidum* n. sp. in the Black-fronted Piping-guan after reintroduction into the wild.

Reintroduced birds face stressors during handling and transportation and when they are released into a novel territory, causing immune suppression that hamper individual responses to parasites (Dickens et al., 2009; Sainsbury and Vaughan-Higgins, 2012). Therefore, such stressors might predispose birds to infection relapses due to activation of exo-erythrocytic merogony with a consequent increase in parasitemia during the chronic stage of infection (Valkiūnas, 2005). This could impair the establishment of infected birds during the period soon after release. In fact, parasitic infections can reduce host survival after release, being an important limiting factor for reintroduction programs (Mathews et al., 2006). Consequently, birds infected by haemosporidians may not be eligible for translocation, as recommended in the conservation program of hihis (*Notiomystis cincta*) in New Zealand (Ewen et al., 2012). On the other hand, Seychelles warblers (*Acrocephalus sechellensis*) infected by *Haemoproteus nucleocondensus* survived after being translocated to new islands in the western Indian Ocean (Fairfield et al., 2016), with some individuals suppressing the infection (Hammers et al., 2016). These contrasting approaches and results show that long-term survival rates should de compared between infected and non-infected Black-fronted Piping-guan after release, to measure the impacts of *H. paraortalidum* n. sp. infection in this reintroduction program.

Preventive treatment against prevalent parasites can be applied before the release of infected birds (Ewen et al., 2012) and this measure can be considered for infected Black-fronted Piping-guans. Single-dose treatment with primaquine, for instance, reduced *Haemoproteus* parasitemia and increased reproductive fitness (Marzal et al., 2005; Merino et al., 2000) and survival (la Puente et al., 2010) in free-living passerines. Benefits on reproductive fitness were also observed in *Plasmodium relictum*-infected birds treated with three doses of Malarone^TM^ (Knowles et al., 2010). Treatment against haemosporidians would be beneficial if those improvements in reproductive fitness and survival were observed in Black-fronted Piping-guan, as the main goal of this reintroduction program is to increase wild populations via reproduction of released birds. Costs of such medications can hamper massive treatment, but efforts should be made to detect and treat infected birds at the pre-release stage to evaluate long-term benefits of this measure.

A parasite haplotype identical to *H. paraortalidum* n. sp. was detected twice in a free-living passerine species (Yellow-breasted Flycatcher *Tolmomyias flaviventris*) captured in an Atlantic forest fragment situated 200 km distant from the reintroduction area of Black-fronted Piping-guan (Lacorte et al., 2013), confirming that this parasite circulate in the wild. These two areas are included in the same biome, but we do not know whether vectors of *H. paraortalidum* n. sp. also occur in the reintroduction area. Same *Haemoproteus* lineages can be transmitted by different species of biting midges (Atkinson, 1991; Bukauskaitė et al., 2015; Žiegytė et al., 2017), increasing the likelihood of *H. paraortalidum* n. sp. being transmitted in the reintroduction area. One next step to the current reintroduction program would be to evaluate haemosporidian diversity in Black-fronted Piping-guan born in the wild, to assess whether *H. paraortalidum* n. sp. is established in this population.

Parasites of subgenus *Parahaemoproteus* commonly have host-ranges restricted to bird order levels, and the same species usually do not complete life cycle in birds belonging to different orders (Valkiūnas, 2005). The detection of *H. paraortalidum* n. sp. in the Yellow-breasted Flycatcher is likely to be a result of abortive infection, probably at sporozoites or exo-erythrocytic stages (Valkiūnas et al., 2014; Valkiūnas and Iezhova, 2017). Moreover, we tested 151 juvenile and adult individuals of Alagoas curassow (*Pauxi mitu*), a Cracidae species managed in CRAX Brazil, but none of them were infected with *H. paraortalidum* n. sp. (unpublished data). This is an intriguing result given that the Alagoas curassow and the Black-fronted Piping-guan are closely related (belong to the same family). Both bird populations share the same environment at the conservation centre, so they are likely to be exposed to the same vectors, as biting midges commonly have broad host ranges (Santiago-Alarcon et al., 2013, 2012). This indicates that *H. paraortalidum* n. sp. may have a restricted range of competent host, although we did not test other Galliformes birds in this facility for the presence of haemosporidians.

Our results demonstrate the importance of sampling numerous individuals belonging to the same species at each study site during different periods of year. This provides opportunities to obtain more infected birds and to collect images of large number of parasites for morphological characterization during light parasitemia. In this study, parasitemia was light and the identification of a new species of *Haemoproteus* was readily possible due to the combined evaluation of gametocytes from different infected individual birds. Microscopic examination and sequencing results from 15 individuals revealed a monoinfection, confirming that parasites from different individuals constitute the same taxonomical unit.

The high prevalence of *H. paraortalidum* n. sp. in a population which members are being reintroduced into the wild raises questions of how to improve the success of reintroduction programs. Information about the impact of *H. paraortalidum* n. sp. infection on the Black-fronted Piping-guan is limited, but it seems that this parasite does not cause major impacts on the established captive population during light chronic infection (Motta et al., 2013). However, we do not know the consequences of *H. paraortalidum* n. sp infection during primary stages of infection and how the parasite behaves in released birds. Treatment of *Haemoproteus* infections remains insufficiently studied (la Puente et al., 2010; Marzal et al., 2005; Merino et al., 2000), but should be recommended at the pre-release stage of birds to evaluate its benefits in reproductive success and in survival of reintroduced individuals.

## Acknowledgements

This work was supported by CoordenaÇão de AperfeiÇoamento de Pessoal de Nível Superior (CAPES), Conselho Nacional de Desenvolvimento Científico e Tecnológico (CNPq), FundaÇão de Amparo à Pesquisa do Estado de Minas Gerais (FAPEMIG). FCFJ was supported by the National Postdoctoral Program/CAPES (PNPD/CAPES). The authors thank the Program for Technological Development in Tools for Health-PDTIS-FIOCRUZ for use of its facilities. The authors thank CRAX Brazil — Wildlife Research Society for allowing sample collection. The funders had no role in study design, data collection and analysis, decision to publish, or preparation of the manuscript.

